# Endothelial cell Piezo1 promotes vascular smooth muscle cell differentiation on large arteries

**DOI:** 10.1101/2024.06.11.598539

**Authors:** Javier Abello, Ying Yin, Yonghui Zhao, Josh Maurer, Jihui Lee, Cherokee Bodell, Abigail J. Clevenger, Zarek Burton, Megan E. Goeckel, Michelle Lin, Stephanie Grainger, Carmen M. Halabi, Shreya A. Raghavan, Rajan Sah, Amber N. Stratman

## Abstract

Vascular stabilization is a mechanosensitive process, in part driven by blood flow. Here, we demonstrate the involvement of the mechanosensitive ion channel, Piezo1, in promoting arterial accumulation of vascular smooth muscle cells (vSMCs) during zebrafish development. Using a series of small molecule antagonists or agonists to temporally regulate Piezo1 activity, we identified a role for the Piezo1 channel in regulating *klf2a* levels and altered targeting of vSMCs between arteries and veins. Increasing Piezo1 activity suppressed *klf2a* and increased vSMC association with the cardinal vein, while inhibition of Piezo1 activity increased *klf2a* levels and decreased vSMC association with arteries. We supported the small molecule data with *in vivo* genetic suppression of *piezo1* and *2* in zebrafish, resulting in loss of *transgelin+* vSMCs on the dorsal aorta. Further, endothelial cell (EC)-specific *Piezo1* knockout in mice was sufficient to decrease vSMC accumulation along the descending dorsal aorta during development, thus phenocopying our zebrafish data, and supporting functional conservation of Piezo1 in mammals. To determine mechanism, we used *in vitro* modeling assays to demonstrate that differential sensing of pulsatile versus laminar flow forces across endothelial cells changes the expression of mural cell differentiation genes. Together, our findings suggest a crucial role for EC Piezo1 in sensing force within large arteries to mediate mural cell differentiation and stabilization of the arterial vasculature.

## Introduction

Blood vessel formation and stabilization are fundamental components of early stage development, supporting intercellular exchange of nutrients, oxygen, and metabolites for organ formation. The mature vascular network—namely arteries, veins, and capillaries—consists of a complex network of highly regulated vessels that vary in size, volume, strength, and elasticity throughout tissues^1–5^. Across the vascular plexus, the vascular wall is composed of three distinct layers—the tunica intima, composed of endothelial cells, the tunica media composed of mural cells (smooth muscle cells and pericytes), and the tunica adventitia mainly composed of fibroblasts, extracellular matrix and progenitor cells^6^. Tight interaction and crosstalk between these different cell types and vessel wall layers is essential for dictating vascular maturation and remodeling^7, 8^.

Mural cells—vascular smooth muscle cells (vSMC) on large vessels and pericytes on capillaries— give blood vessels strength and flexibility in response to hemodynamic forces ^8–11^. vSMC accumulation on arteries is a process that occurs across all vertebrates during the early stages of development in response to mechanical forces. In the zebrafish, this accumulation starts between 2 and 3 days post fertilization (dpf) and follows a rise in pulse pressure after the onset of blood flow ^9, 12–21^. As this relationship between vSMC and pulse pressure occurs in all vertebrates, the current paradigm is that vSMC recruitment happens in response to high shear, pulsatile flow in the arterial vasculature. As veins have uninterrupted straight line flow prior to the formation of venous valves, they fail to recruit vSMCs to the same degree. However, what the molecules are that sense these forces to dictate changes in cellular behavior currently remain unknown.

Kruppel like factor 2a (*Klf2a*) is a blood flow-responsive zinc finger transcription factor that has a wide variety of functions in the developing vasculature. Klf2a expression is regulated in response to differential blood flow forces ^22–27^, and work by our lab and others has demonstrated that throughout development, *klf2a/Klf2* expression is initially observed more highly in veins than arteries in both zebrafish and mouse ^21^. We also demonstrated that *klf2a*-deficient zebrafish mutants accumulate vSMC to the cardinal vein much earlier in development than their wild type counterparts, despite these fish maintaining both *klf2b* and *klf4* expression, suggesting that Klf2a is sufficient to act as a negative regulator of vSMC accumulation^21^. Given that *klf2a* is a transcription factor, there needs to be an upstream force sensor that activates it, and we hypothesized that Piezo channels may be this upstream sensor. In the outflow tract of the zebrafish, this relationship has been previously reported, i.e. that Piezo is a negative regulator of *klf2a* expression ^28^, however, it is still unknown if this is specific to the outflow tract or if it is more generalizable to the vasculature as a whole.

Piezo channels are a family of mechanosensitive, non-selective cation channels expressed in animals, plants, and other eukaryotes, with known effects on vascular development and patterning ^9, 29–37^. In vertebrates, the two members of the Piezo family—Piezo1 and Piezo2—share around 41% sequence homology and are highly conserved across species. Composed of 38 transmembrane domains that form a homotrimer structure arranged in a propeller blade shape, Piezo channels allow for cation entry, such as calcium, into the cell in response to membrane tension ^38, 39^. As such, we sought to understand whether different mechanical forces originating from blood flow patterns would be sufficient to activate the membrane tension sensing role of the Piezo1 channel to regulate *klf2a* and influence the association of vSMCs with the dorsal aorta.

## Results

### Pulsatile blood flow promotes the expression of mural cell differentiation markers

The presiding hypothesis in the field is that differences in blood flow forces between arteries and veins dictate where vSMCs will accumulate and differentiate during development to promote vessel stabilization. Despite this, the molecular pathways regulating vSMC accumulation as a function of mechanosensation are poorly understood. We sought to understand if differences in blood flow patterns between arteries and veins during development alter EC mechanotransduction to regulate directed vSMC motility and differentiation. Zebrafish, like other vertebrate organisms, require blood flow to accumulate *transgelin+* (*tagln*) vascular associated smooth muscle cells along their arteries (vSMCs) during development. In the absence of flow, stopped by the addition of BDM (2,3-Butanedione 2-monoxime, a myosin ATPase inhibitor) to inhibit cardiac muscle contraction ^40^, there is loss of vSMCs (green; *Tg(tagln:NLS-EGFP-2A-CFP-FTASE)^y450^*) associated with blood vessels (magenta; *Tg(kdrl:mCherry-CAAX)^y171^*) (**Fig. 1A-D**).

**Figure 1.**
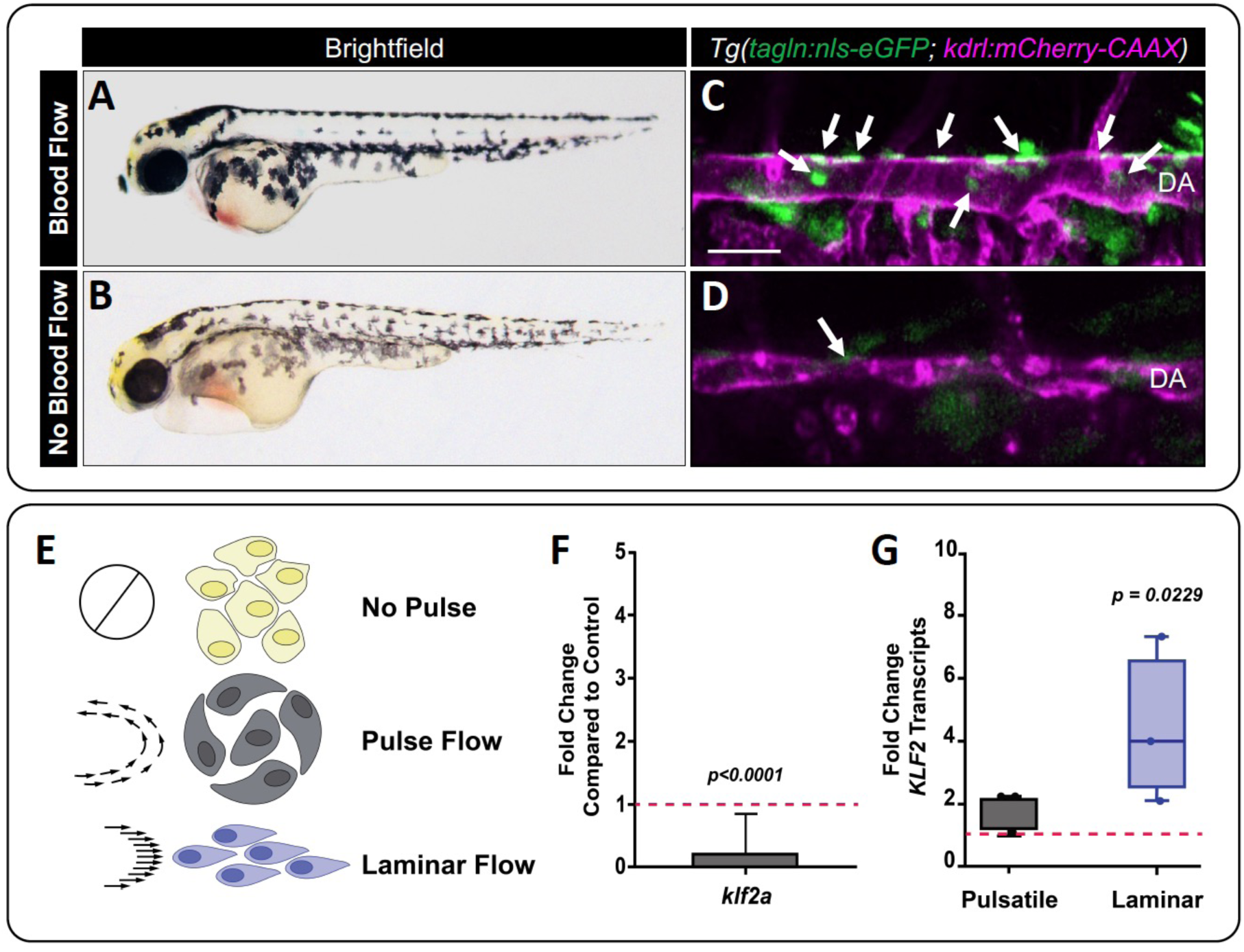
Pulsatile versus continuous laminar flow differentially regulates *klf2* expression. **A,B**) 4 dpf zebrafish embryo with normal blood flow (A) compared to a sibling zebrafish treated with 20 mM of BDM (2,3-butanedione monoxide) at 48 hours to stop the blood flow (B). **C,D**) Confocal image of the medial trunk of a 4 dpf *Tg(tagln:nls-eGFP; kdrl:mCherry-CAAX)* zebrafish with normal blood flow (C) versus those treated with BDM (D). The endothelium is pseudo colored magenta, and *tagln*+ smooth muscle cell nuclei are shown in green and identified by the white arrows. **E**) Schematic representation of no flow, pulsatile flow, and laminar flow conditions. The white arrows inside the rectangular structures indicate the direction of the flow at a given time. **F**) qPCR analysis of *klf2a* transcript levels in 48 hpf zebrafish embryos without blood flow. The dotted line represents *klf2a* expression levels under standard blood flow conditions; (*p = 0.0001*, N= 3 experiments). **G**) *KLF2* transcript levels in HUVECs exposed to pulsatile or continuous laminar flow. The dotted line represents *KLF2* expression under no flow conditions. Expression of *KLF2* in the laminar flow condition was statistically significant compared with no-flow control; (*p = 0.0229*, N= 3 experiments). All statistics were analyzed using the nonparametric Kruskal-Wallis test with Dunn’s multiple comparison test.

Previous reports from our group have shown that the blood flow regulated transcription factor *klf2a* has polarized expression during early embryonic vascular development, with higher expression levels in the venous vasculature over the arterial vasculature in both zebrafish and mice at developmental stages prior to vSMC association ^21^. This expression profile inversely correlates with the level of pulsatile flow going through a vessel (**Video 1**), suggesting that the more ‘pulsation’ that is sensed by the endothelium, the lower *klf2a* transcript levels will be. Developmentally, there has to be blood flow to stimulate *klf2a* transcription (**Fig. 1E,F**, red line denotes the expression level of *klf2a* in animals with blood flow); however, accurately manipulating steady state laminar versus pulsatile flow to study *klf2* expression in the animal is difficult. Therefore, to uniformly apply a type of flow force across a cell population and analyze the KLF2 response, we used human endothelial cells (HUVECs) and *in vitro* modeling, where we can control the parameters of shear stress and pulsation frequency (i.e. set to a human heart rate). From this, we demonstrate that when blood flow is pulsatile, as would be seen in the developing aorta, *KLF2* levels are only modestly upregulated above the no flow condition (dotted line; **Fig. 1E,G**). However, when blood flow is steady state laminar, with no pulsation to it, *KLF2* transcript levels are upregulated 3-7 fold above the no flow condition (**Fig. 1E,G**). This suggests that ECs regulate *KLF2/klf2a* expression in response to the type of fluid flow within the developing vasculature, with laminar steady state flow able to upregulate *KLF2* to a greater extent than pulsatile flow.

We then modified this *in vitro* model to test the hypothesis that EC flow sensing altered vascular mural cell behavior and differentiation (**Fig. 2A**). GFP^+^ human brain vascular pericytes (HBVPs) were selected for these studies because they express lower levels of mural cell differentiation genes than contractile, differentiated vSMCs do, allowing us to determine if less differentiated mural cell populations could gain differentiation markers in response to different types of flow forces. GFP-HBVPs were seeded below a monolayer of ECs with a thin layer of collagen type I between the two cell populations, and either pulsatile or laminar flow applied across the EC monolayer. Using time-lapse live cell imaging, we found that pericytes moved a greater total distance in response to pulsatile flow compared to laminar flow across the ECs (**Fig. 2B,C; Videos 2 and 3**). Pericytes in the pulsatile flow assays also had altered aspect ratios after 24 hours compared to pericytes in laminar flow assays (**Fig. 2D; Videos 2 and 3**). Assessing mural cell differentiation markers in response to laminar versus pulsatile flow, we saw modest changes in *PDGFRB* transcript or PDGFRB protein levels between the two conditions (**Fig. 2E,H**), but saw a significant increase in both *ACTA2* and *TAGLN* transcripts and ACTA2 and TAGLN protein levels in response to pulsatile flow compared to laminar flow (**Fig. 2F,G,I-K**). Taken together, these findings suggest that there is a mechanosensitive protein in the endothelium that differentially senses pulsatile (i.e. arterial) versus laminar (i.e. venous) forces (**Video 1**) to control mural cell differentiation.

**Figure 2.**
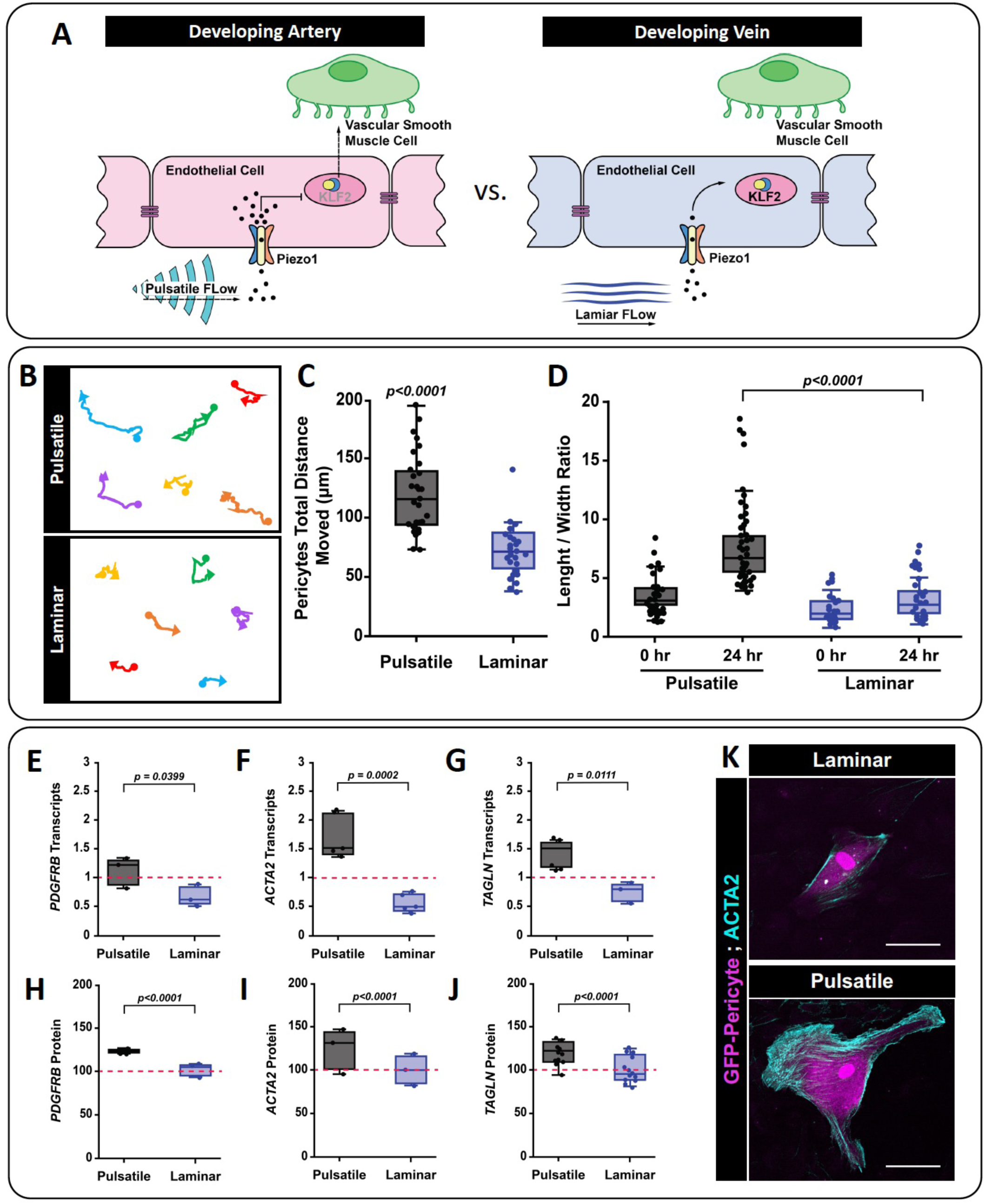
Blood flow type influences mural cell differentiation *in vitro*. **A**) The schematic representation shows the proposed relationship between blood flow types, mechanical forces, and different patterns of vSMC association that occur between arteries and veins. We hypothesize that Piezo1 is the mechanosensor responsible for sensing hemodynamic forces, playing a role in vSMC association and differentiation. DA: dorsal aorta; scale bar = 50 um. **B**) Representative tracking of HBVP-GFP movements under laminar versus pulsatile flow. The filled circle marks the start of the track and the arrowhead at the end. **C**) Quantification of HBVP-GFP total movement under pulsatile (N=38) or continuous laminar (N=33) flow. Unpaired t-test (*p=<0.0001*). **D**) Average HBVP-GFP length-to-width aspect ratios at 0 hours or after 24 hours under pulsatile versus laminar flow. Statistical analysis was performed using Kruskal-Wallis test with Dunn’s multiple comparison test (*p= <0.0001*, N=63). **E-G**) qPCR transcript levels of mural cell differentiation markers after 24 hours of laminar or pulsatile flow—(E) platelet-derived growth factor receptor beta *PDFGRB* (*p=<0.0399,* uncorrected Fisher’s LSD; N=3); (F) alpha-smooth muscle actin (*ACTA2*) (*p=0.0002,* Tukey’s multiple comparison test; N=5); and (G) Transgelin (*TAGLN*) (*p=<0.0111*, Tukey’s multiple comparison test; N=3). All data is normalized to GAPDH. **H-J**) Quantification of relative protein levels of mural cell differentiation markers via immunostaining analysis after 24 hours of laminar or pulsatile flow—(H) platelet-derived growth factor receptor beta PDFGRB (*p=<0.0001,* Mann-Whitney two-tail t-test; N=3); (I) alpha-smooth muscle actin (ACTA2) (*p=<0.0001,* Mann-Whitney two-tail t-test; N=5); and (J) Transgelin (TAGLN) (*p=<0.0001,* Mann-Whitney two-tail t-test; N=3). **K**) Representative images of ACTA2 immunostaining following HBVP-GFP exposure to different flow types for 24 hours. scale bar = 50 um.

We sought to understand if Piezo1 in ECs could serve as this force sensor using the well characterized pharmacologic agonists—Yoda1 and Jedi2—and the antagonist—GsMTx4—to activate or suppress Piezo1 activity in an acute and temporal manner (**Supp. Fig. 1A**). To ensure that these small molecules could impact Piezo channel activity in the zebrafish, we used *Tg(actb2:GCaMP6s)^stl351^*transgenic zebrafish to live image relative Ca^2+^ concentrations in the endothelium and surrounding cells following treatments of interest (**Supp. Fig. 1A,B**). We found that embryos treated with 50 nM GsMTx4 showed a decrease in relative Ca^2+^ concentration, while embryos treated with 400 nM Jedi2 showed an increase in relative Ca^2+^ concentration (**Supp. Fig. 1C,D**), demonstrating that these small molecules are functional in the zebrafish. To assess the impact of Piezo1 functional modulation on blood flow sensing, zebrafish embryos were treated with these compounds and assessed for *klf2a* levels. Consistent with previous literature ^28^, we note a modest reciprocal relationship between Piezo1 activity and *klf2a* expression—where increasing Piezo1 activity via Jedi2 or Yoda1 led to a decrease in *klf2a* transcript levels (in zebrafish; **Supp. Fig. 2A,B**), while inhibition of Piezo1 activity via GsMTx4 led to increased *klf2a* transcript levels (in zebrafish; **Supp. Fig. 2B**). Concurrently, this relationship between Piezo1 and KLF2 was also validated in human ECs by examining human KLF2 protein (**Supp. Fig. 2C**). Together, these data suggest that Piezo1 can regulate *klf2a*/KLF2 expression based on its activity and provide a link between blood flow, mechanotransduction, and the endothelium.

### Piezo1 regulates *transgelin* (*tagln*) positive vSMC association with large arteries during development

To test the requirement of Piezo channels for mural cell, particularly vSMC, recruitment and/or differentiation, we used a multi-species approach. Suppressing Piezo activity in the zebrafish starting at 24 hpf using GsMTx4 led to a decrease in the number of *tagln*+ vSMCs associated with the dorsal aorta with no changes in the number of vSMCs associated with the cardinal vein (**Fig. 3A-C**). To complement these data, we analyzed *piezo1^sa12608^/piezo2a.1^sa12414^* genetic zebrafish mutants (**Fig. 3D-F**). Double homozygous *piezo1^-/-^/piezo2a.1^-/-^* mutants developed marked edema, displayed a disrupted body axis, and had decreased blood flow, impacting our ability to interpret phenotypes from this condition. However, analysis of the dorsal aorta versus the cardinal vein of *piezo1^+/-^/piezo2a.1^-/-^*mutants and *piezo1^-/-^/piezo2a.1^+/-^* mutants showed a decreased number of *tagln*+ vSMCs associated with the dorsal aorta (**Fig. 3D-F**), consistent with the GsMTx4 pharmacologic data. Importantly, these zebrafish mutants had normal blood flow, supporting a role for *piezo1/piezo2a.1* in vSMC differentiation along the axial vasculature.

**Figure 3.**
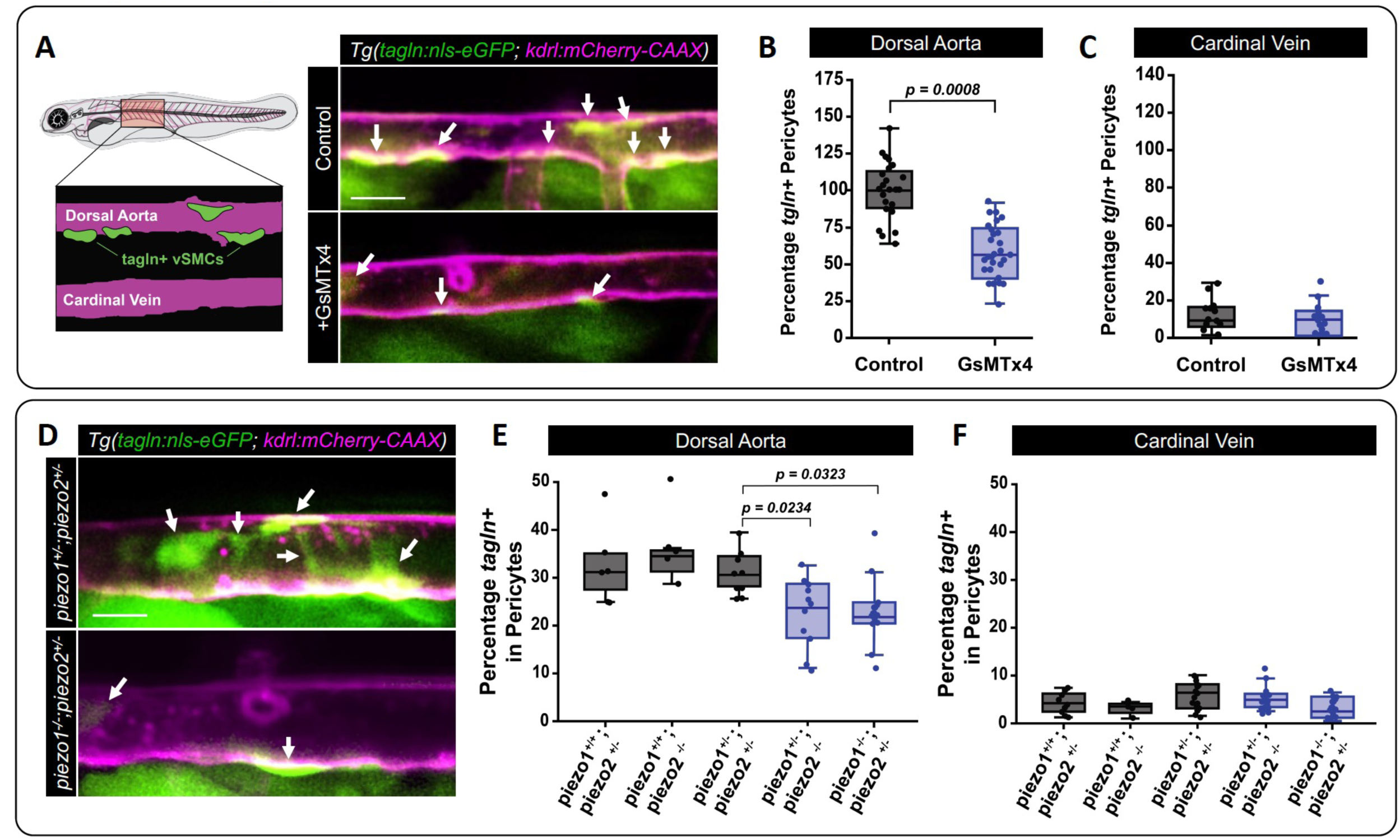
Inhibition of Piezo1 impairs vSMC association with the dorsal aorta in zebrafish. **A**) Schematic of a 96 hpf zebrafish embryo and the imaging area collected within the zebrafish for analysis. Confocal images of the medial trunk of *Tg(tagln:eGFP; kdrl:mCherry-CAAX)* zebrafish treated with 50 nM of GsMTx4 or water as the vehicle control. vSMCs shown in green and the endothelium in magenta. Arrows highlight vSMCs associated with the dorsal aorta. **B,C**) Quantification of *tagln* positive vSMCs associated with (B) the dorsal aorta or (C) the cardinal vein following treatment with GsMTx4 (*p=<0.0008*, unpaired two-tailed t-test; N=25 and 30). **D**) Confocal images of the medial trunk of *Tg(tagln:eGFP; kdrl:mCherry-CAAX)* zebrafish carrying mutations to *piezo1* and *piezo2a.1*. Arrows highlight vSMCs associated with the dorsal aorta. **E,F**) *piezo1^+/-^;piezo2a.1^-/-^* individuals were crossed to *piezo1^+/-^;piezo2a.1^+/-^* individuals to generate the indicated genotypes. Quantification is shown of *tagln* positive vSMCs associated with the dorsal aorta (E) or cardinal vein (F) *piezo1^+/-^; piezo2^+/-^* (N=10); *piezo1^+/+^; piezo2^+/-^* (N=6); *piezo1^+/+^; piezo2^-/-^* (N= 4); *piezo1^+/-^; piezo2^-/-^* (N=10); *piezo1^-/-^; piezo2^+/-^* (N=10). *piezo1^+/-^; piezo2^+/-^* vs. *piezo1^+/-^; piezo2^-/-^* (*p=0.0234*), *piezo1^+/-^; piezo2^+/-^* vs. *piezo1^-/-^; piezo2^+/-^* (*p=0.0323*). The box plots show the median versus the first and third quartiles of the data. The whiskers indicate the spread of data within 1.5x above and below the interquartile range. Statistics were preformed using one-way ANOVA with Tukey’s multiple comparisons test. *piezo1^-/-^;piezo2a.1^-/-^*double mutants lack robust blood flow and have edema; therefore, they were excluded from the analysis. scale bar = 50 um.

Conversely, treatment with either Piezo1 activator—Yoda1 or Jedi2—led to a modest increase in the number of vSMCs on the dorsal aorta but a significant increase in the number of vSMCs on the cardinal vein (**Fig. 4A-C**). We were able to rescue the Yoda1 effects using its antagonist, Dooku (**Supp. Fig 3**), supporting specificity of Yoda1 in targeting Piezo1 in these functions. The increase in vSMCs on the cardinal vein when Piezo1 is activated was consistent with our previous studies demonstrating that genetic suppression of *klf2a* levels in zebrafish leads to increased differentiation of vSMCs on veins and our current findings that increasing Piezo activity can suppress *klf2a*/Klf2 transcript and protein levels in some contexts (**Supp. Fig. 2**). We did not note changes in vessel patterning, as addition of the small molecules started after the initial vascular plexus was formed (**Supp. Fig. 4A-C**), or zebrafish heart rate (**Supp. Fig. 4D,E**) following these treatments. Together, these data suggest that Piezo channel activity, and likely Piezo1 more specifically, regulate vSMC localization to the vasculature.

**Figure 4.**
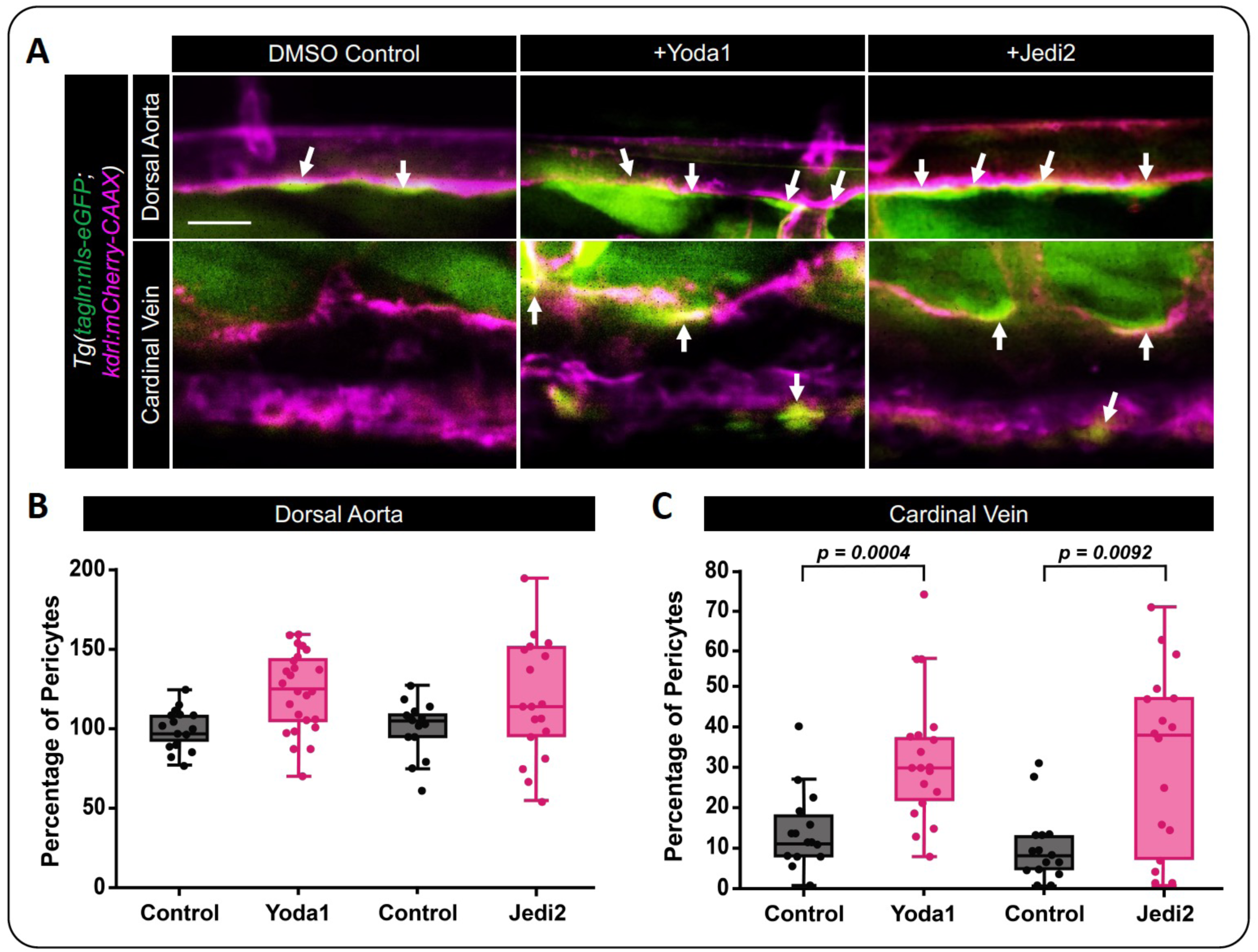
Activation of Piezo1 activity induces vSMC’s association with the cardinal vein. **A**) Confocal images of the medial trunk of *Tg(tagln:eGFP; kdrl:mCherry-CAAX)* zebrafish treated with 10 nM of Yoda1, 400 nM of Jedi2 or DMSO as the vehicle control. vSMCs shown in green and the endothelium in magenta. Arrows highlight vSMCs associated with the dorsal aorta and cardinal vein. **B,C**) Quantification of *tagln* positive vSMCs associated with (B) the dorsal aorta or (C) the cardinal vein following treatment with Yoda1, Jedi2, or DMSO (control). DMSO vs Yoda1 (*p=0.0004*; N=21 and 25); DMSO vs Jedi2 (*p=0.0092*; N=15 and 18). Statistical analysis was performed using a Kruskal-Wallis test with Dunn’s multiple comparison test. Scale bar = 50 um.

Next, we generated EC-specific *Piezo1* knockout mice (*Piezo1^ecko^*) where exons 20-23 of Piezo1 were flanked by loxP sites^41^ and excised using an endothelial-specific Cre driver, *VE-Cad:Cre*^42^, to assess the EC specific requirements for Piezo1. At E12.5-E13.5, *Piezo1^ecko^* animals appeared to be paler than *Piezo1^flox/flox^*siblings (**Fig. 5A**), likely consistent with previously reported phenotypes of vessel loss following Piezo1 loss of function in the vasculature ^31–33^. To assess vSMC association with the large axial vessels, we sectioned these embryos followed by immunostaining with endomucin (vessels, green) and αSMA (vSMCs, magenta). *Piezo1^ecko^* animals had reduced vSMC wall thickness around the descending dorsal aorta and an approximately 50% reduction in the number of αSMA+ vSMC layers that make up the vessel wall (**Fig. 5B-D**). These data support the conclusion that EC Piezo1 is critical for modifying the environment needed to support vSMC differentiation along the aorta.

**Figure 5.**
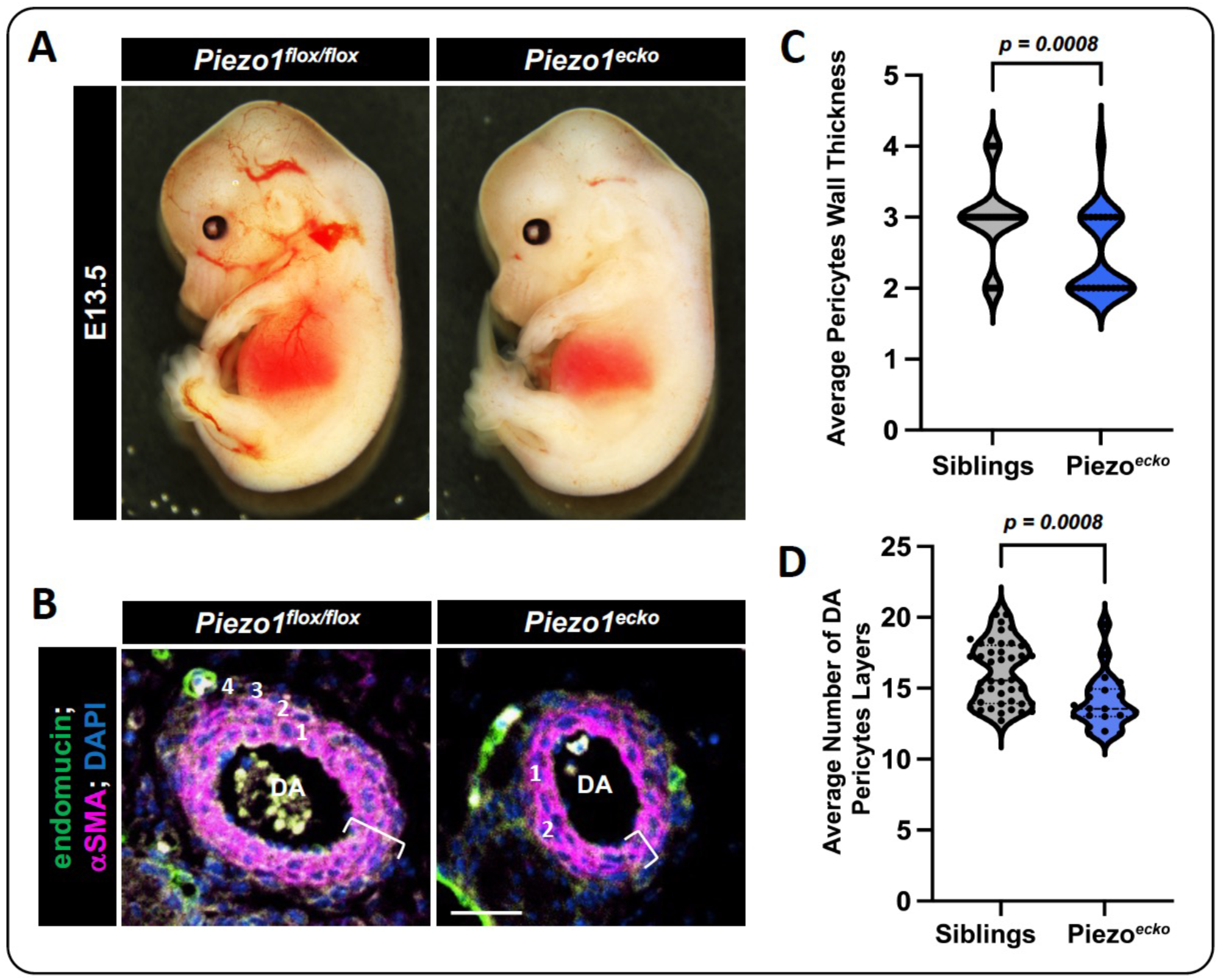
EC-specific Piezo1 knockout reduces vSMC association with the dorsal aorta in mice. **A**) *VE-Cad:Cre* mice were crossed to *Piezo1^flox//flox^* mice to generate E12.5-13.5 embryos across genotypes. Representative images of a *Piezo1^flox//flox^* sibling and an EC-specific Piezo1 deficient embryo (*Piezo1^ecko^*) are shown. **B**) Immunostaining of cross sections of the descending dorsal aorta (DA) in *Piezo1^flox/flox^* versus *Piezo1^ecko^* E12.5-13.5 embryos. The endothelium is labeled with endomucin in green, vSMCs are labeled with alpha-smooth muscle actin (⍺SMA) in magenta, and nuclei (DAPI) are shown in blue. The white bracket indicates the wall thickness of the the vSMC layer, the white numbers indicate the number of vSMC layers present. **C**) Quantification of the average vSMC wall thickness (um) in *Piezo1^flox/flox^* and *Piezo1^ecko^* embryos (*p=0.0008*, N=42 siblings; N=20 *Piezo1^ecko^*). Statistical analysis was performed using a two-tail unpaired t-test. **D**) Quantification of the average number of vMSC layers on the descending DA in *Piezo1^flox/flox^*and *Piezo1^ecko^* embryos (*p=0.0008*, N=42 siblings; N=20 *Piezo1^ecko^*). Statistical analysis was performed using a two-tail unpaired t-test. Scale bar (B) = 50 um.

As our data strongly predict a role for EC Piezo1 mechano-sensing in altering localized signaling and mural cell activity, we assessed the role of Piezo1 in regulating EC-pericyte interactions using *in vitro* live imaging assays. In these assays, the cells were encapsulated in a 2.5 mg/ml, concave topped, 3D collagen gel with no perfusion or flow associated with the culture. Therefore, any forces sensed by cells in this encapsulated system would be strain fields generated as a function of hydrogel encapsulation, contributing cell-level compressive strains in the absence of externally applied forces. The cells were added and mixed together in a 1:5 pericyte to EC ratio in the gel and then allowed to self-assemble over 72 hours into lumen containing EC tubes with pericytes associated along the tube (**Fig. 6A**). In the control condition, approximately half of the pericytes in the culture associated with an EC tube in this time frame, while half did not (**Fig. 6B**, showing pericytes on the EC tube). Adding Jedi2 to the cultures increased the number of pericytes associated with EC tubes to approximately 70% (**Fig. 6B**), while GsMTx4 treatment reduced the number of pericytes associated with an EC tube to approximately 25-30% (**Fig. 6B**). We next carried out EC specific siRNA treatments and found that suppression of EC Piezo1 (EC siPiezo1) led to impaired association of pericytes with EC tubes compared to cultures that used EC siControl cells (**Fig. 6C,D**). The pericytes were wild type and untreated with any modifiers across these siRNA assays, suggesting the signals dictating their association were altered in the endothelium specifically. Given that we could alter Piezo1 activation above and below control levels in this assay, this could imply that the primary force that Piezo1 senses from blood flow is a compression force.

**Figure 6.**
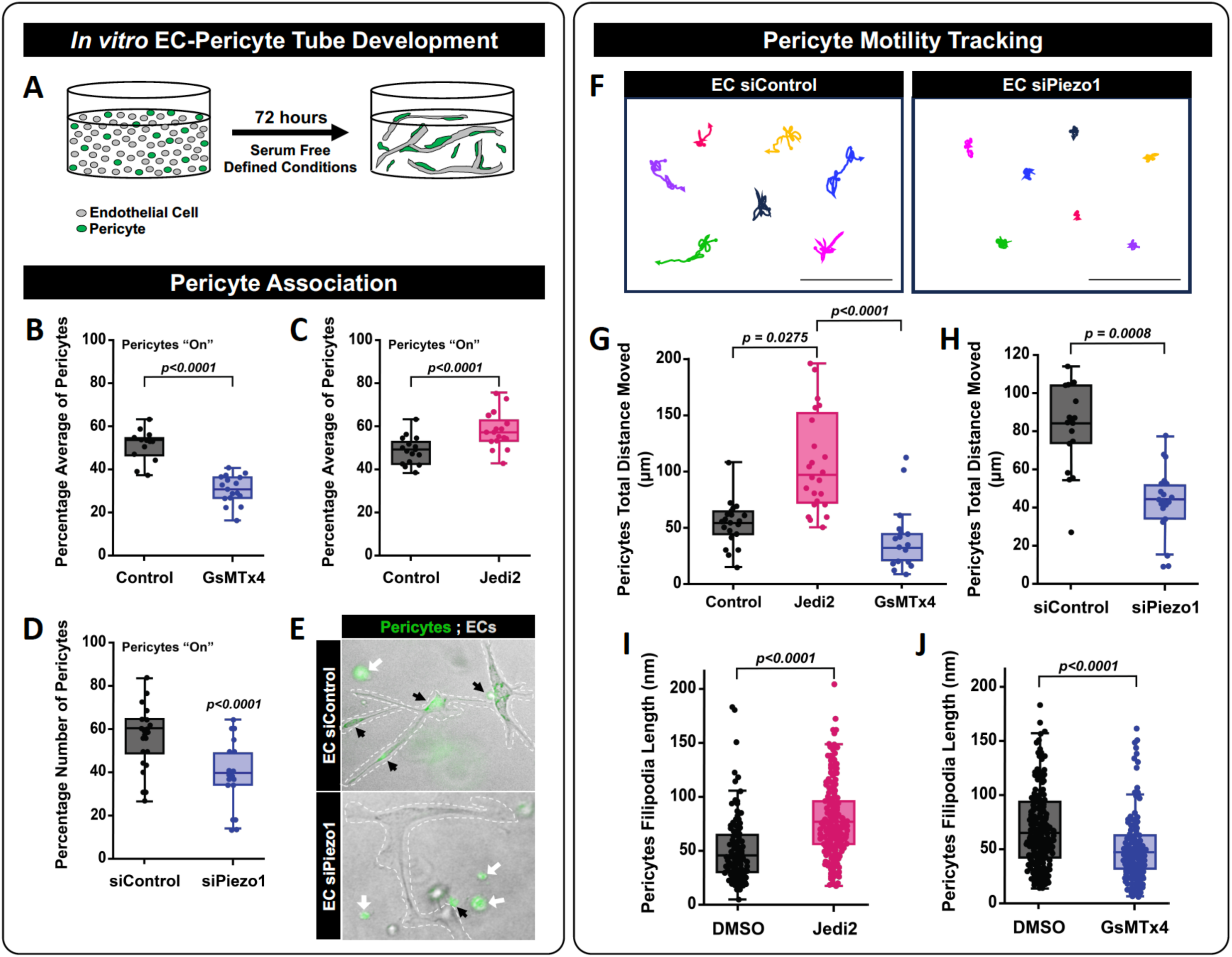
3D *in vitro* modeling of endothelial cell and pericyte interactions can be regulated by Piezo1 activity. **A**) Schematic representation of our 3D cell culture model. HUVECs (EC) and GFP-HBVPs (pericyte) were seeded in a 5:1 ratio, respectively, in a 3D collagen type I gel and allowed to self-assemble for 72 hours. Piezo1 activity was modulated pharmacologically by treating the cell cultures with 200nM Jedi2 or 50nM of GsMTx4 versus DMSO (control) or by siRNA suppression of EC Piezo1 (siPiezo1) versus an siRNA control (siControl). After 72 hours of incubation, pericyte association with EC tubes was evaluated. **B**) Quantification of pericyte colocalization with EC tubes under treatments with control (water) and GsMTx4. (*p=<0.0001*, N= 15 independent cultures). Statistical analysis was performed using a two-tail unpaired t-test. **C**) Quantification of pericyte colocalization with EC tubes under treatments with control (DMSO) and Jedi2. (*p=<0.0001*, N=3-5 independent cultures). Statistical analysis was performed using a two-tail unpaired t-test. **D)** Quantification of pericyte colocalization with EC tubes following EC specific treatment with Piezo1 siRNA (*p=<0.0001*, N=30 independent cultures). Statistical analysis was performed using a two-tail unpaired t-test. **E**) Representative images of EC/GFP-pericyte cocultures. EC tubes are outlined by the white dashed line, and GFP-pericytes are green in siControl and EC siPiezo1 conditions. **F**) Representative cell tracking plots of GFP-pericytes in 3D cocultures over 48 hours. **G,H**) Quantification of GFP-pericyte total motility in 3D cocultures, obtained from 48-hour time-lapse video tracking of cultures treated with (H) Jedi2 (*p=0.00275*) or GsMTx4 (N=12-25 independent cells); or (G) EC siPiezo1 cultures (*p<0.0001*, N=24 independent cells). Statistical analysis was performed using one way ANOVA with a Tukey’s multiple comparison test. **I,J**) Quantification of GFP-pericyte filipodia length in 3D cocultures (I) DMSO vs Jedi2 (*p<0.0001*, N=200 cells), (J) DMSO vs. GsMTx4 (*p<0.0001*, N=200 cells). Phenotypes were assessed after 72 hours in Jedi2 (I) GsMTx4 (J). Statistical analysis was performed using a two-tail Mann-Whtney test. Scale bar = 50 um.

To determine if changes in pericyte localization were due to altered motility, we used time-lapse microscopy to track pericyte motility across these assays. We found that following Jedi2 treatment, pericytes moved more distance over time than pericytes in control cultures. Conversely, GsMTx4 or EC siPiezo1 treated cells led to a suppression in pericytes motility compared to their counterpart control condition (**Fig. 6E-G**). Consistent with their motility capacity, cultures treated with Jedi2 had pericytes that exhibited longer filipodia (**Fig. 6H**) and cultures treated with GsMTx4 had pericytes that exhibited shorter filipodia (**Fig. 6I**). Together these data support a hypothesis that EC Piezo1 can likely be activated by outward, compression forces from pulsatile shear on ECs themselves to direct mural cell motility and differentiation across time and alter vascular stabilization during development.

## Discussion

Understanding how blood flow, mechanics, and force sensing interface to alter mural cell (i.e. both vSMCs and pericytes) differentiation and/or recruitment to vessels remains an active area of investigation for the field. Deciphering how blood vessels are built and stabilized across development is critical for the diagnosis and treatment of a number of congenital vascular disorders, such as congenital heart defects (CHD), which can manifest as developmental abnormalities of the great vessels (aorta and cardinal vein). In fact, several studies have demonstrated the role of Piezo1 in CHD and the development of vascular valves. Continued work deciphering the molecular and cellular pathways controlling these defects remains of utmost importance for truly understanding these pathologies. Here, we describe a unique role for the EC Piezo1 activity in regulating vSMC differentiation, to stabilize the developing vascular network (**Fig. 3-5**).

Cumulatively, our data supports the conclusion that Piezo1 activation in the endothelium helps dictate the production of paracrine factors that alter vSMC differentiation during development. The expression and activation of Piezo1 correlate with repression of Klf2 transcript and protein levels, and in previous work we have demonstrated that Klf2 can suppress the expression of vSMC pro-differentiation and proliferative signals such as PDGFB ^21^. However, loss of PDGFB/PDGFRB function in vSMCs on large vessels is not fully penetrant like it is in other contexts such as the brain, leaving open the question of what downstream signaling pathways are additionally activated to contribute to vSMC recruitment and differentiation on the dorsal aorta. Further, whether the link between Piezo1 and Klf2 is causative of altering downstream gene programs or just correlative in response to blood flow sensing will be a focus of future studies.

One of the difficulties of studying a mechanosensitive protein such as Piezo1 *in vivo*, is the inability to decouple what specific forces are being sensed in the context of genetic loss of function. As such, we have utilized novel tools *in vitro* to begin to address this issue (**Fig. 2 and 6**). From these studies, we conclude that pulsatile shear stress helps promote the expression of mural differentiation markers, and that strain, or membrane stress on ECs in 3D without flow present, is sufficient to reveal roles for Piezo1 in regulating EC-mural cell interactions. This sets up the interesting hypothesis that Piezo1 might predominantly sense the outward pulsatile force (strain) propagated through developing large arteries from the beating heart, rather than shear itself ^15^. While this is a compelling hypothesis, the platforms to test these relationships are still being developed by the field, but will allow for exciting new findings moving forward.

## Materials and Methods

### Zebrafish Lines Husbandry

The work with Zebrafish (*Danio rerio*) and the experimental procedures developed and implemented in this study were reviewed and approved by the Washington University in St. Louis School of Medicine Institutional Animal Care and Use Committee (IACUC). *piezo1^sa12608^* (ZL9041.03) and *piezo2a.1^sa12414^*(ZL8936.10) mutants were crossed with the transgenic vascular background *Tg(tagln:nls-eGFP-2a-CFP-FTASE ^y450Tg^/kdrl:mCherry-CAAX ^y171Tg^)*^15^ to allow imaging of transgelin positive vSMC on vessels. To track Piezo1 activity, we utilize the *Tg(actb2:GCaMP6s^stl351^)* ^43^. The *piezo1* and *piezo2a.1* mutations utilized in this study are Single Nucleotide Polymorphisms (SNPs). To genotype these fish lines, we implemented the Kompetitive Allele Specific PCR assay or KASP. The KASP assay for *piezo1* and *piezo2a.1* were synthesized by LGC Biosearch Technologies under project number 1166.002. To genotype the adult zebrafish, the fish were anesthetized with 100mg/mL of MS-222 (Syndel USA, cat. #: NC0872873) for less than 2 minutes to cut a small piece of the tail fin for gDNA extraction. gDNA extraction was carried out in 0.2mL 8-Strip PCR tubes (Alkaly Scientific, Cat.#: PC7061), and gDNA extraction buffer (10mM Tris-HCl pH: 8.2, 200mM NaCl, 0.5% SDS, and 200μg/mL of Proteinase K) at room temperature. The KASP reaction was prepared following the manufacturer’s recommendations in 1X Master mix (Bioserch Technologies, Cat. #: KBS-1050), and the alleles amplification was performed using a QuantStudio 3 Real-Time PCR Systems (ThermoFisher Scientific, Cat.#: A28567).

### Cell culture

Human Umbilical Vein Endothelial Cells (HUVEC) (ATCC, Cat. #: PCS100010) were culture in 1X M-199 Medium (Gibco, Cat. #: 12340030) supplemented with 16 % FBS (Gibco, Cat. #: 16140071), 15mg Endothelial Cell Growth Supplement (Corning, Cat. #: 35400), 50mg of Heparin sodium salt (Sigma-Aldrich, Cat. #: H3393) and 1X Antibiotic-Antimycotic (Gibco, Cat. #: 15240096). HUVEC were seeded at an initial concentration of 3000 cell/cm^2^ in gelatin-coated (1mg/mL in PBS) flasks. Cells were incubated at 37°C, 5% CO_2_, and 95% humidity. GFP labeled Human Brain Vascular Pericytes (HBVP) (ScienceCell™, Cat. #: 1200) were grown in DMEM (Gibco, Cat. # 11995065) supplemented with 10% FBS and 1X Antibiotic-Antimycotic under the cell culture conditions indicated above. HBVP cells were transfected with lentiviral particles (GenTarget, Cat.#: LVP444-G) to express GFP following the manufacturer’s recommendations, in brief: HBVP cells were seeded at an initial concentration of 1.3X10^4^ cells/cm^2^ in T-175 cell culture flasks (ThermoFisher Scientific, Cat.#: 178883) pre-coated with gelatin (4mg/mL in 1X PBS) and incubated overnight at 37°C, 5% CO_2_, and 95% humidity. Then 1X10^7^ IFU viral particles were added to the cell cultures and incubated for an additional 72 hours. After transfection, the cells were observed under a fluorescent microscope to detect the GFP signal. Then, the cell cultures were treated with 2μg/mL of puromycin (Sigma-Aldrich, Cat.#: P8833) to select GFP-positive HBVP cells.

### Pharmacological modulation of Piezo channel

To modulate Piezo activity *in vivo* and *in vitro,* we treated the experimental units with the Piezo1 inhibitor GsMTx4 (Tocris, Cat. #: 4912) at a concentration of 50nM; and the Piezo1 activators Yoda1 (Tocris, Cat. #: 5586) at 10nM and Jedi2 (Tocris, Cat. #: 6614) at 400nM. We inhibit Yoda1 activity with Dooku1 (Tocris, Cat. #: 6568) at 10nM. The activity of each compound was evaluated against their suspension vehicle as control: water for GsMTx4 and DMSO for Yoda1 and Jedi2. The zebrafish embryos were treated with the indicated dose of each compound at 24 hpf, and the fish water containing the small molecules was replaced every 24 hours. The embryos were visualized at 96 hpf following. Data was acquired using a Nikon Ti2E microscope equipped with a CSU-W1 spinning disk confocal (Yokogawa).

### Blood Flow Stop and TRAP assay

24 hpf zebrafish embryos were treated with 20mM of the inhibitor 2,3-Butanedione 2-monoxime (BDM) (Sigma-Aldrich, Cat. #: B0753). Translation ribosomal affinity purification (TRAP) at 48 hpf. After 24 hours of treatment, the embryos were treated with Deyolking buffer (55mM NaCl, 1.9mM KCl, 1.25mM NaHCO3), supplemented with 1X Proteinase inhibitor (Sigma-Aldrich, Cat. #: 8340). Then the embryos were rinsed with deyolking wash buffer (10mM Tris-HCl at pH 8.5, 110mM NaCl, 3.5mM KCl, 2.7mM CaCl_2_) before proceeding with the maceration of the tissue in homogenization buffer (50mM Tris-HCL at pH7.4, 100mM KCl, 12mM MgCl_2_, 1% NP-40) supplemented with: 1mM DTT Sigma-Aldrich, Cat.#: 646563), 200 U/mL RNasin (Promega Cat.#: N2111), 100 ug/mL cycloheximide (Sigma-Aldrich, Cat.#: 7698) and 1mg/mL of heparin (Sigma-Aldrich, Cat.#: H3393). To capture the ribosomes first, the samples were incubated for 16 hours at 4°C with HA Tag-precoated Dynabeads-Protein G (Invitrogen, Cat.#: 10007D). Then the Dynabeads were rinsed with a high salt buffer (50mM Tris pH 7.4, 300mM KCl, 12mM MgCl_2_, 1% NP-40, 1 mM DTT, 1X protease inhibitors, 200 units/mL RNAsin, 100ug/mL cycloheximide, 1mg/mL heparin) and the ribosomal bounded-RNA was extracted with the Total RNA Mini Kit IBD (IBI Scientific, Cat. #: IB47302).

### Flow assay

The flow experiments were conducted as previously described ^44^, in brief:1.3X10^5^ HBVP-GFP cells were seeded on a gelatin pre-coated µ-Slide ^0.4^I Luer chambers (Ibibi^®^, Cat. #: 80176). After 16 hours of incubation, we created a layer of collagen type I (Corning, Cat.#: 354249) by diluting 100μg/mL of collagen in EC-complete medium. We allow polymerization of the layer by incubating the cell cultures for 30 minutes at 37°C, 5% CO_2_, and 95% humidity. When the collagen layer was formed, the remaining of the collagen-EC medium solution was aspirated and 2.5X10^5^ HUVEC cells per slide were seeded on top of the collagen. The HBVP/HUVEC co-culture was incubated overnight under standard tissue culture conditions. The next morning, the slides were transferred onto the flow system (Flocel, Cat. #: WPX1) adapted to produce pulsatile flow between 12-15 dyn/cm^2^ at 60 RPM or continuous laminar flow at 15 dyn/cm^2^ to resemble physiological conditions^44^. Complete EC culture medium was flowed across the cells for 24 hours. At the termination of experiments, the cultures were rinsed 3 times with 1X PBS and either lysed for DNA/mRNA isolation or fixed with 4% PFA for 10 minutes or 99.9% Methanol for 5 minutes, depending on the downstream application.

### Immunofluorescence staining of paraffin sections and flow slides

Mouse embryos were embedded in paraffin for sectioning. The slides were washed 3 times for 15 minutes in HistoChoice (Sigma-Aldrich, Cat.#: H2779), followed by consecutive 15 15-minute washes in 100%, 95%, and 75% ethanol, and a final wash in 1x PBS. To retrieve the antigen, the slides were submerged in 10mM Sodium Citrate Buffer and placed in a pressurized chamber at 100°C for 20 minutes. The slides were then treated with 1X Tris Glycine for 15 minutes at room temperature, followed by 0.1% Triton X-100 for 30 minutes. After the permeabilization step, the slides were incubated for 1 hour at room temperature in blocking buffer (20mM Tris-Base, 150mM NaCl, and 0.1% Tween 20 at pH: 7.6) supplemented with 5% of specific serum or Bovine Serum Albumin (BSA), depending on manufacturer’s recommendation. Antibodies: Endomucin V.7C7 antibody (Santa Cruz Biotechnology, Cat.#: sc-65495) at 1:50 dilution in blocking buffer 5% donkey serum, followed by a secondary FITC Goat Anti-Rat Ig, (BD-Pharmigen, Cat #: 554016) at 1:500 dilution in 1X PBST 0.2% BSA; α-Smooth Muscle Actin-Cy3, mouse monoclonal antibody (Sigma-Aldrich, Cat.#: C-6118) at 1:500 dilution in in blocking buffer 5% donkey serum; Hoechst 34580 (Sigma-Aldrich, Cat.#: 63493) at 1:5000 dilution in blocking buffer 5% donkey serum.

Immunostaining of flow assays followed the same general protocol outlined above, without rehydration washes and antigen retrieval. Antibodies: α-Smooth Muscle Actin (D4K9N) Rabbit monoclonal antibody (Cell Signaling, Cat.#: 19245) at 1:100 dilution in dilution in blocking buffer 1% BSA, followed by the secondary antibody Alexa Fluor 633 goat anti-rabbit IgG (Invitrogen, Cat.#: A21071) at 1:2000 dilution in blocking buffer 1% BSA.

### Western Blot

Protein was obtained from HUVEC cells treated with the Piezo1 pharmacological modulators. Cells monolayers were lysed in RIPA lysis buffer (ThermoFisher Scientific, Cat.#: 89901) supplemented with phosphatase inhibitor (ThermoFisher Scientific, Cat.#: A32957) and protease inhibitor (ThermoFisher Scientific, Cat.#: A32955). The cell lysate was collected and centrifuged at 14000 xg, 4°C for 15 minutes to eliminate insoluble proteins. The lysate protein content was quantified with the Pierce™ BCA Protein Assay Kits (ThermoFisher Scientific, Cat.#: 23225) following the manufacturer’s recommendations. Proteins were denatured at 85°C for 2 minutes, and electrophoresis was carried out in NOVEX™ Tis-Glycine gels (ThermoFisher Scientific, Cat.#: XP04205BOX) at 125V, 100 minutes. After electrophoresis, the proteins were transferred to PVDM membrane using the dry transferring system iBlot 2 (Invitrogen). The membrane was blocked overnight in 1X TBST 5% nonfat dry milk; then incubated for 1 hour at room temperature with Klf2 anti-rabbit (Aviva Systems Biology Corporation, cat. #: ARP32760_P050) 1:500 dilution, followed by mouse anti-rabbit IgG-HRP (Santa Cruz Biotechnology, Cat.#: sc-2357) 1:1000 dilution. After images were captured, the Klf2 antibody was stripped using the Restore™ Western Blot Stripping Buffer (ThermoScientific, Cat. #: 21059) for 15 minutes at room temperature. GAPDH anti-goat IgG (Abcam, Cat. #: AF5718) 1:5000 dilution, followed by secondary anti-goat HRP (R&D systems, Cat.#: HAF019) 1:5000 dilution was then blotted to as a loading control.

### 3D Collagen Matrix Assays and siRNA treatment

HUVEC and GFP-HBVP cells were grown and maintained following the standard procedures described in the cell culture section. Cells in T-25 flasks were transfected using siPORTAmine (ThermoFisher Scientific, Cat.#: AM4503) and 50nM of siRNA to Piezo1 (Life Technologies, Cat.#: 4390827) versus a negative control 2 siRNA (Life Technologies, Cat.#: 4390847). Following 3 days of incubation after the last siRNA treatment, cells were trypsinized and collected for use in the 3D collagen matrix assays. siRNA-treated HUVECs and GFP-HBVPs were seeded into the collagen I gel using 96-well A/2 plates as was previously described ^45–47^. This serum-free hydrogel contains 2.5 mg/mL Collagen I from rat tail (Corning, Cat.#: 354249), FGF (R&D systems, Cat.#: 233-FB-025/CF), SCF (R&D systems, Cat.#: 255-SC-010/CF), IL3 (R&D systems, Cat.#: 203-IL-010/CF), SDF1a (Cat.#: 350-NS-010/CF) in the gel and ascorbic acid (AA), FGF (R&D systems, Cat.#: 233-FB-025/CF), and IGF-2 (R&D systems Cat.#: 292-G2-250) in the culture media. After 72 hours of incubation at 37°C, 5% CO_2_, and 95% humidity, the cell cultures were fixed in 4% PFA for 30 minutes, and images were acquired with the light microscope EVOS M5000 (Invitrogen).

### Time Lapse and Live Imaging Analysis

Images from zebrafish and immunostaining analysis were acquired using a Nikon Ti2E microscope equipped with a CSU-W1 spinning disk confocal (Yokogawa). Fluorescent Images of the vasculature were taken at the medial trunk of 2-5 dpf zebrafish embryos using a 20x or 40x Plan-Apo objective. For imaging, zebrafish embryos were anesthetized with MS-222 (Syndel USA, cat. #: NC0872873) and mounted in 35 mm optical bottom petri dishes (MatTek Life Science, Cat.#: P35-1.5-20-C) with 0.1% low melt agarose (IBI Scientific, Cat. #: IB70056).

For live cell tracking, images were acquired every 20 minutes for 24-48 hours (depending on the experiment) in an EVOS M7000 microscope (ThermoFisher Scientific) equipped with an on-stage incubator to provide standard cell culture conditions at 5% CO_2,_ 95% humidity, and 37°C.

### GCaMP Imaging and Analysis

*Tg(actb2:GCaMP6s)^stl351^* transgenic zebrafish were incrossed and allowed to develop for 48 hours. At 48 hpf, embryos were mounted in 1% low melt point agarose and allowed to acclimate in fish water for 30 min. Imaging dishes containing the mounted embryos were placed on a Nikon Ti2E microscope equipped with a CSU-W1 spinning disk confocal (Yokogawa). Control images of individual animals were acquired, taking an image every 10 seconds for 5 minutes to establish a baseline of Ca^2+^ activity in each animal. The small molecules were then added to the water in the imaging dish and images acquired of the same animal again (every 10 seconds for 5 minutes) to establish a response in Ca^2+^ activity to the small molecules. Using ImageJ, changes in signal intensity at 5 sites from each animal were measured for changes in Ca2+ flux, in 4 independent animals. This data was then Fourier transformed, using the function in Excel, to normalize oscillating patterns across this data and plotted for visualization.

### Mouse husbandry and genotyping

To obtain the vascular endothelial-specific Piezo1 knockout mice for this study, the strain B6.Cg-Piezo1*^tm2.1Apat^*/J^41^ (The Jackson Laboratory, stock #: 029213) was crossed with the VE-cadherin-Cre strain B6.Cg-Tg(Cdh5-cre)1Spe/J^42^ (The Jackson Laboratory, stock #033055). To obtain timed embryos, pregnant females were euthanized with CO_2_, embryos were removed, rinsed, and fixed with 4% PFA. The yolk sacks and a piece of the tail from the adult female were collected for DNA extraction and genotyping. Genotyping was carried out by touchdown PCR as follows: Piezo1 wild type was amplified with the primers, P1_F: 5’-CTT GAC CTG TCC CCT TCC CCA TCA AG-3’ AND P1_R: 5’-CAG TCA CTG CTC TTA ACC ATT GAG CCA TCT C-3’. Piezo1 knockout was amplified with the primers, P1KO_F: 5’-CTT GAC CTG TCC CCT TCC CCA TCA AG-3’ and P1KO_R: 5’-AGG TTG CAG GGT GGC ATG GCT CTT TTT-3’, initial denaturation at 94°C for 3 minutes followed by 10 cycles at 94°C for 20 seconds, 65°C for 15 seconds with a reduction of 0.5°C per cycle, 68°C for 40 seconds followed by 28 cycles at 94°C for 15 seconds, 62.5°C for 15 seconds, 72°C for 15 seconds, followed by a final extension at 72°C for 2 minutes. The VE-cadherin-Cre was amplified with the primer, CDH5-Cre_F: 5’-AGG CAG CTC ACA AAG GAA CAA T-3’ and CDH5-Cre_R: 5’-TCG TG CAT CGA CCG GTA A-3’, initial denaturation at 94°C for 2 minutes followed by 10 cycles at 94°C for 20 seconds, 65°C for 15 seconds with a reduction of 0.5°C per cycle, 68°C for 10 seconds followed by 28 cycles at 94°C for 15 seconds, 60°C for 15 seconds, 72°C for 10 seconds, followed by a final extension at 72°C for 1 minute. The animals were maintained in the animal facility of the School of Medicine at Washington University in St. Louis under standard husbandry conditions, including social housing, a 12-hour light-dark cycle, 23°C, water *ad libitum,* and fed with regular chow food.

### qPCR

In this study, RNA was extracted from adult zebrafish tail fins, embryos, HUVEC, and HBVP cells using a total RNA isolation kit (IBI Scientific, Cat.#: IB47302) following the manufacturer’s recommendations. 1ug of mRNA was converted to cDNA using the SuperScript™ IV VILO™ Master Mix kit (Invitrogen, Cat.#: 11766050). Gene expression was performed using Taqman™ probes for each gene of interest. HBVP cell differentiation was evaluated with the probes: PDGFRB Hs01019589_m1 (ThermoFisher Scientific, Cat.#: 4331182), TAGLN Hs01038777_g1 (ThermoFisher Scientific, Cat.#: 4331182) and ACTA2 Hs00426835_g1 (ThermoFisher Scientific, Cat.#: 4331182). KLF2 was evaluated in human cells with the probe Hs07291763_gH (ThermoFisher Scientific, Cat.#: 4331182) and in zebrafish with *klf2a*: Dr03138475_g1 (ThermoFisher Scientific, Cat.#: 4331182). The transcript levels were normalized to GAPDH with human Hs99999905_g1. (ThermoFisher Scientific, Cat.#: 4331182) or zebrafish Dr034336842_m1 (ThermoFisher Scientific, Cat.#: 4331182). RT-qPCR was carried out using the TaqMan Fast Advance master Mix (ThermoFisher Scientific, Cat.#: 4444557) in a QuantStudio3 qPCR machine (ThermoFisher Scientific, Cat.#: A28131)

### Statistics

Statistical analysis was conducted with the software GraphPad Prism v10.0. For two-variable comparisons under normality assumption with the Shapiro-Wilk test, the p-values were determined with the two-tailed, unpaired t-test. In the case of data that was not following normal distribution, we analyzed it with a nonparametric Mann-Whitney U test. For more than two variables, a two-way ANOVA test was employed with the Kruskal-Wallis test for nonparametric data and Uncorrected Fisher’s LSD test or Dunn’s test for multiple comparisons. Statistically significant was set at a p-value of less than 0.05. Data graphics were created with the same software.

## Author Contributions

JA, YY, YZ, JM, JL, CB, AC, ZB, MG, ML, SG, CH, SR, RS and ANS conceptualized and performed experiments; JA, YY, JL, CB, and ANS analyzed results and made the figures; JA, SG, CH, SR, RS and ANS designed the research and wrote the paper. Conflict-of-interest disclosure: The authors declare no competing financial interests. Correspondence: Amber N. Stratman, Department of Cell Biology and Physiology, Washington University School of Medicine St. Louis, MO, 63110;

## Acknowledgements

The authors would like to thank members of the Stratman laboratory for their critical comments on this manuscript. NIH/NIGMS R35 GM137976 (A.N.S.); Children’s Discovery Institute of Washington University and St. Louis Children’s Hospital (A.N.S.); *the Washington University Institute of Clinical and Translational Sciences, which is, in part, supported by the NIH/National Center for Advancing Translational Sciences (NCATS), CTSA grant #UL1TR002345* (A.N.S.); Cancer Prevention and Research Institute of Texas RP230204 (S.A.R.); NIH/NIGMS R35 GM142779 (S.G.). The content is solely the responsibility of the authors and does not necessarily represent the official views of the National Institutes of Health.

---

**Supplemental Figure 1.**
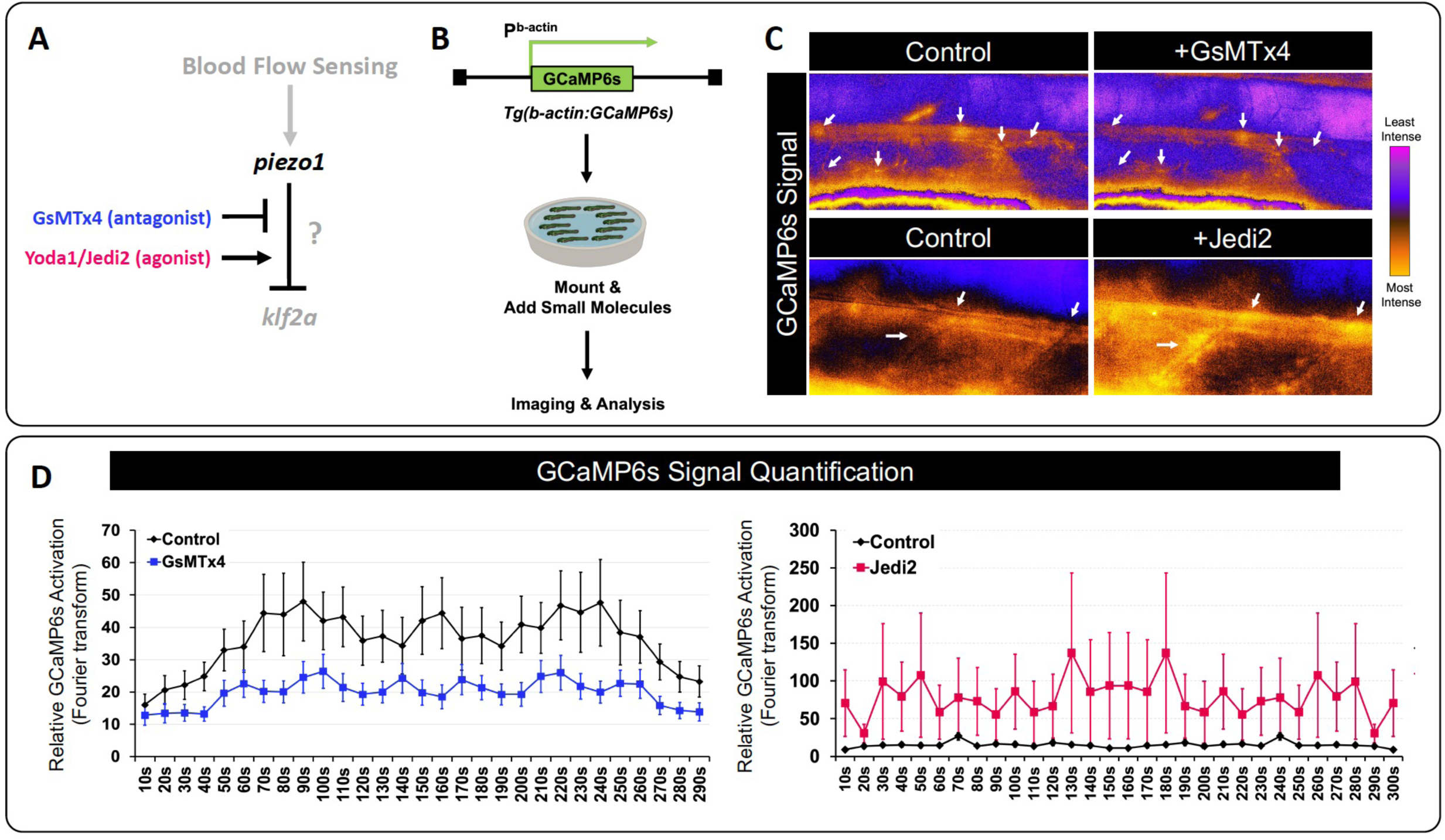
Piezo antagonists and agonists regulate Ca2^+^ flux in the zebrafish. **A**) Schematic of the proposed role of Piezo1 in blood flow sensing, and the available antagonists and agonists to regulate its activity. **B**) Schematic of our imaging set up of *Tg(bact2:GCaMP6s)* embryos for tracking relative Ca^2+^ concentrations in the zebrafish embryo. **C**) Representative images of the GCaMP6s signal pseudocolored for ease of viewing. Orange represents sites of highest signal intensity, while magenta represents sites of lowest signal intensity. The control panels indicate the GCaMP6s signal in response to DMSO addition, followed by images of the same region’s response to the addition of the indicated small molecule. White arrows indicate regions where the GCaMP6s signal is markedly changed. **D**) To quantify the relative changes in Ca^2+^ concentration, individual cells were measured for their GCaMP6s signal over a 5-minute period. Data from all cells was then Fourier transformed and plotted as a relationship between the control and indicated small molecule treatment (GsMTx4 on the left; Jedi2 on the right). N = 4 individual fish; 5 cells regions from each fish, for a total of 20 cells per condition.

**Supplemental Figure 2.**
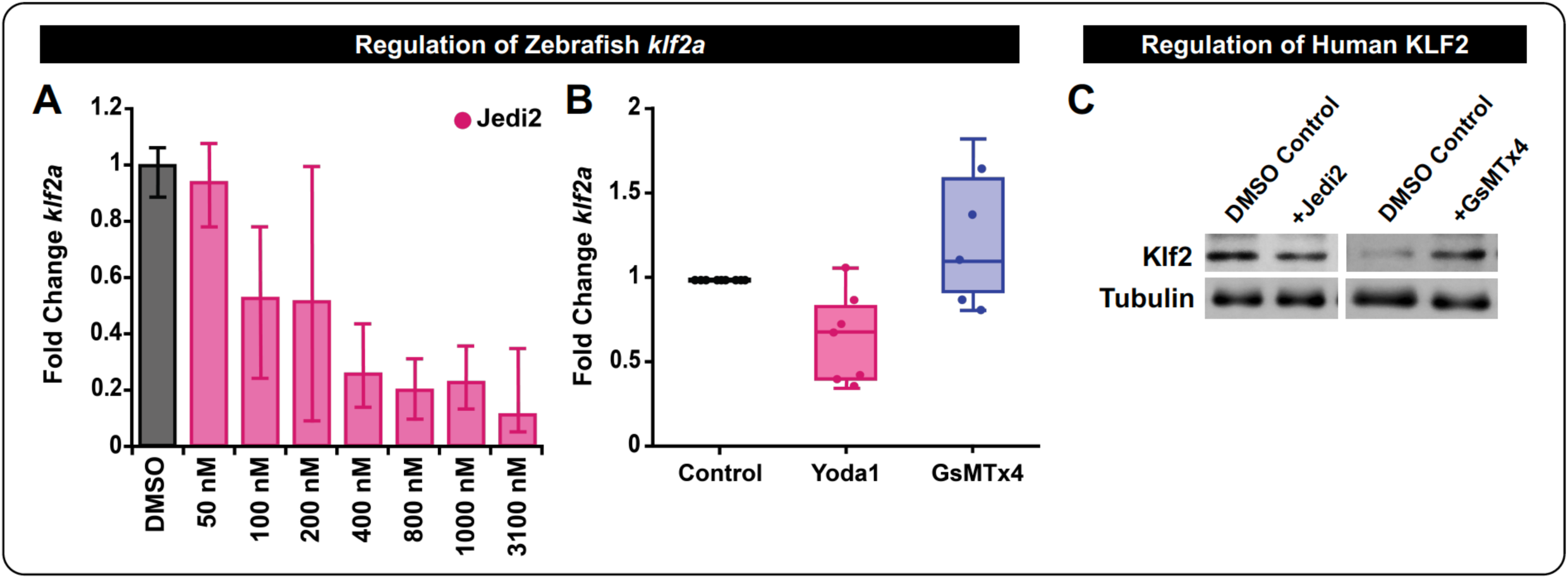
Piezo1 activity regulates *klf2a*/KLF2 levels. **A**) Zebrafish embryos were treated with a dose response curve to Jedi2, starting at 24 hpf. mRNA was extracted at 48 hpf and converted to cDNA for RT-qPCR analysis of *klf2a* transcript levels. Data is presented as a fold change compared to the DMSO treated control siblings. **B**) RT-qPCR analysis for *klf2a* transcript levels in zebrafish embryos treated with 10 nM Yoda1 or 50nM GsMTx4 from 24-48 hpf. Data is normalized to the DMSO control condition. Each dot represents an individual biological replicate: Yoda 1 (N=7), GsMTx4 (N=5). **C**) Western blot analysis of KLF2 protein levels in HUVECs treated with Jedi2 or GsMTx4 for 16 hours, compared to tubulin loading controls. Representative of N=3 biological replicates.

**Supplemental Figure 3.**
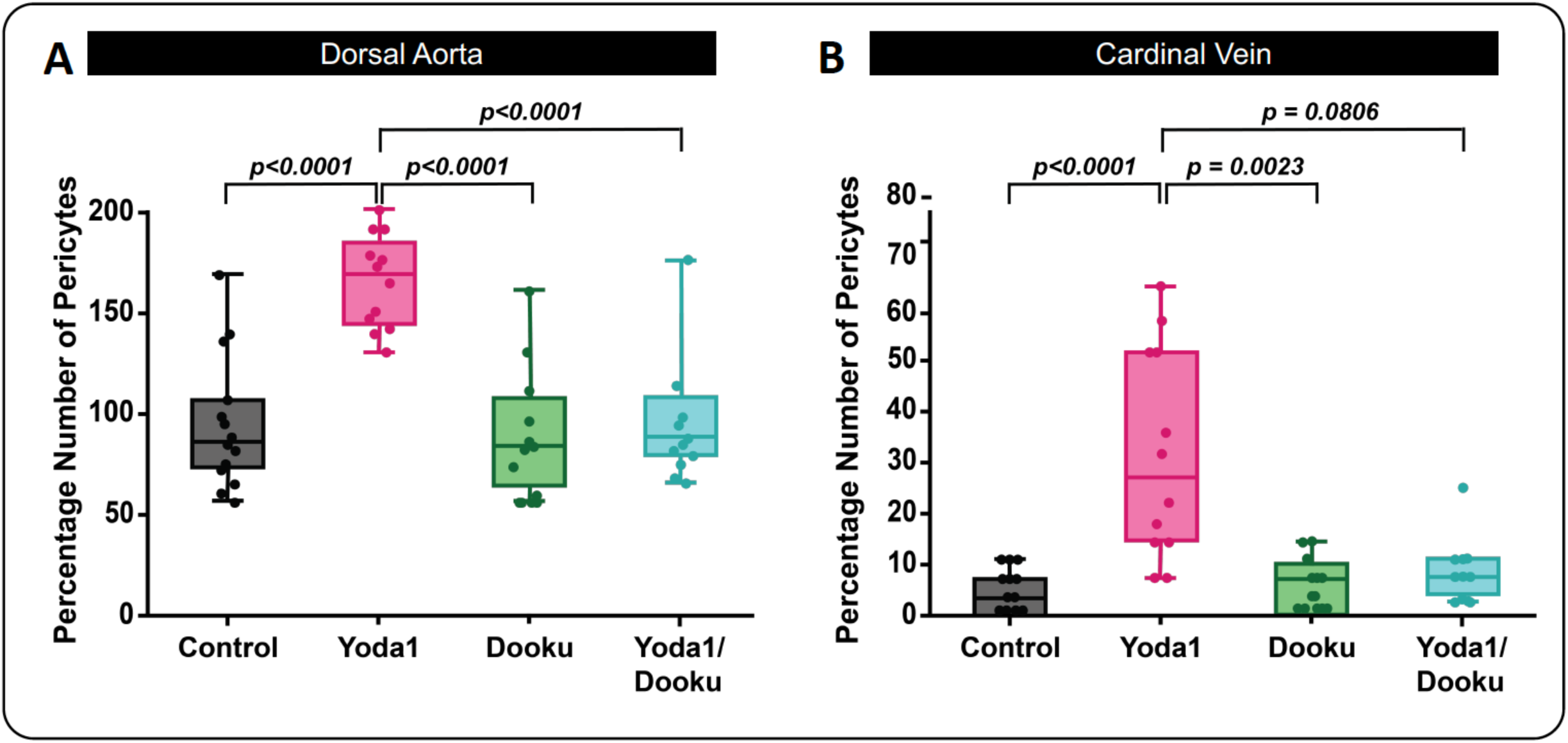
Dooku rescues Yoda1 induced vSMC phenotypes in the zebrafish. **A,B**) Quantification of *tagln* positive vSMCs associated with the dorsal aorta (A) or cardinal vein (B) in zebrafish treated with 10 nM of Yoda1 (N=12); 10 nM of Dooku (N=11), or both (N=11) versus DMSO vehicle control (N=14), starting at 24 hpf and analyzed at 96 hpf. Statistical analysis was performed using a Kruskal-Wallis test with Dunn’s multiple comparison test.

**Supplemental Figure 4.**
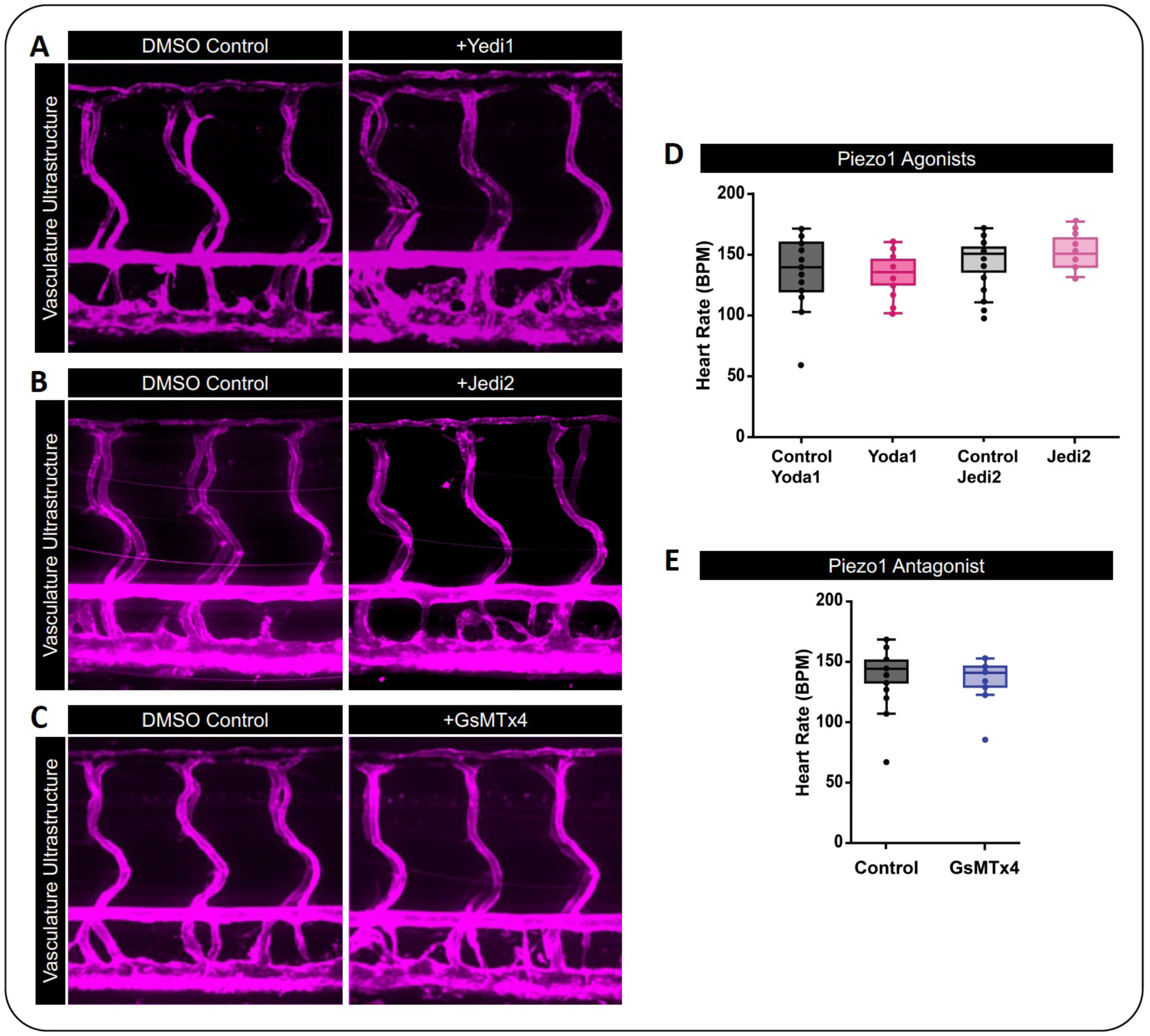
Treatment of zebrafish with Piezo1 small molecules from 24-96 hpf does not alter vascular patterning or heart rate. **A-C**) Confocal spinning disk images from *Tg(kdrl:mCherry-CAAX)^y171^*zebrafish embryos treated with the indicated small molecule versus its respective control. Treatment started at 24 hpf, and imaging was done at 96 hpf. **D,E**) Quantification of zebrafish heart rate (in beats per minute (BMP)) following treatment with Yoda1/Jedi2 (D) or GsMTx4 (E) versus their paired control.

